# Human induced pluripotent stem cell-derived microglia with a CX3CR1-V249I genetic variant exhibit dysfunctional phenotypes and modulate neuronal growth and function

**DOI:** 10.1101/2025.07.01.662163

**Authors:** Kaylee D. Tutrow, Jade Harkin, Laurna Varghese, Melody Hernandez, Kang-Chieh Huang, Yue Fang, Parker Wilcox, Logan Bedford, Tsai-Yu Lin, Chi Zhang, Cátia Gomes, Shweta Puntambekar, Stephanie Bissel, Bruce T. Lamb, Jason S. Meyer

## Abstract

The involvement of microglia in neurodegenerative diseases has drawn increasing attention, as many genetic risk factors are preferentially expressed in microglia. Microglial fractalkine receptor (CX3CR1) signaling regulates many key microglial functions, and the CX3CR1-V249I single nucleotide polymorphism (SNP) has been associated with increased risk for multiple neurodegenerative conditions, including Alzheimer’s disease, yet its functional consequences in human microglia remain unexplored. In this study, we generated iPSC-derived human microglia-like cells (hMGLs) and found that the CX3CR1-V249I variant increased susceptibility to starvation-induced cell death, reduced amyloid-beta uptake, altered microglial morphology, and impaired migration, with more pronounced effects in homozygous cells. Co-culture with neurons demonstrated that hMGLs with the CX3CR1-V249I variant misregulated neuronal properties, including abnormal neuronal growth as well as an induction of neuronal hyperexcitability. These findings highlight the critical role of CX3CR1 in regulating microglial function and implicate the V249I variant in driving pathogenic microglial states relevant to neurodegeneration.

## INTRODUCTION

Microglia, the resident immune cells of the central nervous system (CNS), are essential for maintaining brain homeostasis, clearing toxic substances, supporting neuronal health, and responding to environmental challenges (Del Rio-Hortega Bereciartu, 2020; Ginhoux et al., 2013; Kettenmann et al., 2013). In response to disease, microglia undergo well-characterized phenotypic and transcriptional changes and are increasingly recognized as critical contributors to neurodegenerative processes (Colonna and Butovsky, 2017; Gao et al., 2023; Heneka et al., 2025). Genetic variants expressed in microglia also influence susceptibility to neurodegenerative diseases (Andrews et al., 2020; Bertram and Tanzi, 2008; McQuade et al., 2020; Tsai et al., 2023). As a result, the contributions of microglia to neurodegenerative pathological features have become an important focus of recent research.

The CX3CR1 receptor is selectively expressed on microglia in the central nervous system, and its ligand fractalkine is produced and presented primarily by neurons. The CX3CL1–CX3CR1 signaling axis represents a key mode of communication between neurons and microglia, critically modulating microglial inflammatory responses (Arnoux and Audinat, 2015; Jones et al., 2010). Disruption of this signaling pathway leads to aberrant microglial phenotypes, including impaired phagocytosis and defective migration (Castro-Sanchez et al., 2019; Lastres-Becker et al., 2014). Altered expression and signaling of CX3CR1 has been observed in disease states and is associated with exacerbated neurodegeneration (Bhaskar et al., 2010; Castro-Sanchez *et al*., 2019; Puntambekar et al., 2022). Moreover, polymorphisms in the CX3CR1 gene have been implicated in both peripheral and neurodegenerative diseases (Lopez-Lopez et al., 2018; Stojkovic et al., 2012). In particular, the CX3CR1-V249I SNP reduces receptor function by decreasing the number and affinity of fractalkine binding sites (Moatti et al., 2001) and has been associated with increased risk and severity of Alzheimer’s disease pathology (Lopez-Lopez *et al*., 2018). Although prior studies primarily in rodent knockout models have demonstrated the effects of disrupted fractalkine signaling on AD-related microglial phenotypes, the mechanisms underlying these changes in human systems remain unexplored.

To investigate the functional consequences of the CX3CR1-V249I variant in human microglia, we used human induced pluripotent stem cells (iPSCs) engineered by CRISPR/Cas9 genome editing to generate iPSC lines harboring heterozygous and homozygous CX3CR1-V249I alleles, alongside matched isogenic controls. These iPSC lines were differentiated into human microglia-like cells (hMGLs), enabling assessment of variant-specific phenotypes. CX3CR1-V249I hMGLs exhibited pronounced alterations in proliferation, susceptibility to stress-induced cell death, morphology, and phagocytic capacity. Transcriptomic profiling by RNA-seq revealed that both heterozygous and homozygous lines shared differential expression patterns converging on innate immune signaling pathways. Co-culture of microglia with neurons augmented some neuronal phenotypes, notably enhanced neurite outgrowth and neuronal hyperexcitability. These findings underscore the functional impact of the CX3CR1-V249I variant on human microglia and their interactions with neurons, offering insight into potential mechanisms of neurodegenerative disease susceptibility.

## RESULTS

### CRISPR/Cas9 gene editing to generate isogenic control and CX3CR1-V249I iPSCs

To determine the effects of the CX3CR1-V249I single nucleotide polymorphism (SNP) on human microglia-like cells, we leveraged existing CRISPR/Cas9 approaches to generate isogenic control and CX3CR1-V249I cell lines. A homology-directed repair (HDR) template was designed to include the V429I (c.839C>T) variant, with an additional silent mutation (c.828G>A) (Figure 1A-B) that disrupted the PAM site on the HDR template, helping to ensure specificity of editing to the genomic DNA. Additionally, the introduction of the c.828G>A silent mutation disrupted an AclI restriction site, enabling facile screening of gene editing via restriction digest assays (Figure 1C). A plasmid containing the CX3CR1-V249I HDR template as well as the guide RNA was co-transfected with pCas9-GFP, with the GFP signal used to sort for presumptively edited cells (Figures 1D). Clonal populations were screened by PCR amplification of the edited region and enzymatic digestion with AcII, with prospectively edited clones further sequenced to ensure proper editing of the target gene and identification of both heterozygous and homozygous clones alongside an isogenic control cell line (Figure 1E-F). Additionally, edited clones were screened for likely off-target effects to ensure specificity of editing, with no off-target edits identified at any of the top loci (Supplemental Figure 1).

**Figure 1.**
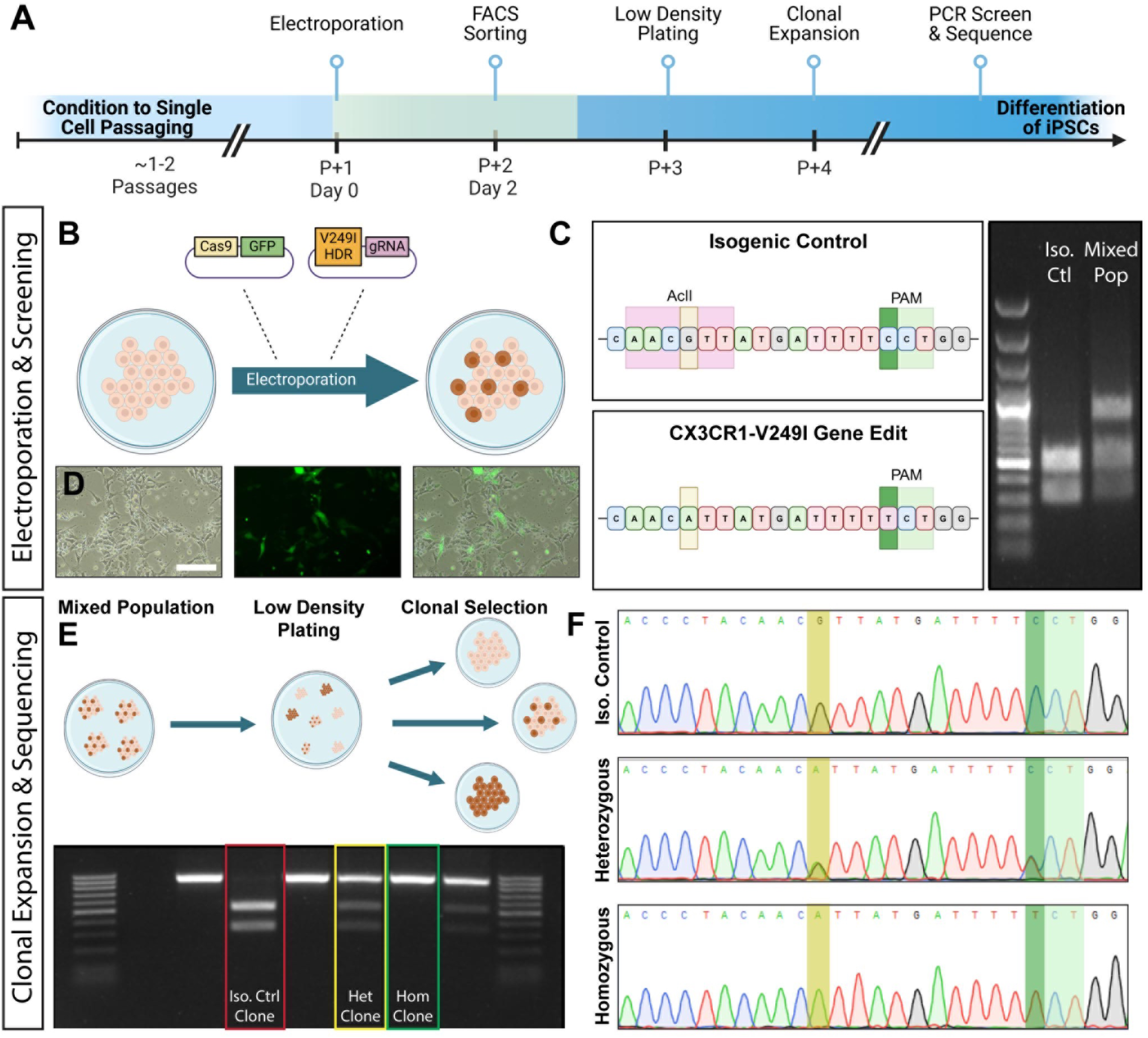
Utilization of CRISPR/Cas9 for development of CX3CR1-V249I cell lines. (A) An overview of the cell editing strategy for the development of CX3CR1-V249I bearing cell lines alongside isogenic control. (B) A wild-type cell line was electroporated with Cas9-GFP plasmid and a homology directed repair template and guide RNA plasmid for the V249I locus. (C) The population was screened for the anticipated removal of the AclI restriction digest site. (D) Transient expression of GFP allowed for FACS sorting with higher efficiency cell selection. (E) Cells from the mixed population were clonally expanded. Clones were screened for edits by restriction digest. (F) Sanger sequencing was used to confirm genotypes. Scale bar: 200 µm.

### Differentiation of human microglial-like cells (hMGLs) from CX3CR1-V249I and isogenic controls

To determine if the CX3CR1-V249I variant disrupts microglial phenotypes, we first sought to differentiate MGLs from iPSCs to determine if differentiation capacity was altered as well as if any morphological or other phenotypic changes resulted from this gene variant. Heterozygous and homozygous CX3CR1-V249I SNP cell lines, alongside the respective isogenic control, were differentiated first into hematopoietic progenitor cells (HPCs) (Supplemental Figure 2), which were then harvested and further differentiated into hMGLs. Immunocytochemical analyses demonstrated that differentiated MGLs across all genotypes expressed microglial markers IBA1, TREM2 and P2RY12, and developed more ramified morphologies indicative of maturation (Figure 2A-I). While no differences in the two-dimensional area were observed dependent upon genotype, both heterozygous and homozygous V249I hMGLs demonstrated an increased cell perimeter and decreased cell circularity compared to controls (Figure 2J-L). Moreover, homozygous V249I hMGLs exhibited an increased number of branches compared to both control and heterozygous V249I hMGLs (Figure 2M), with these morphological differences suggesting the possibility of functional changes due to the V249I variant.

**Figure 2.**
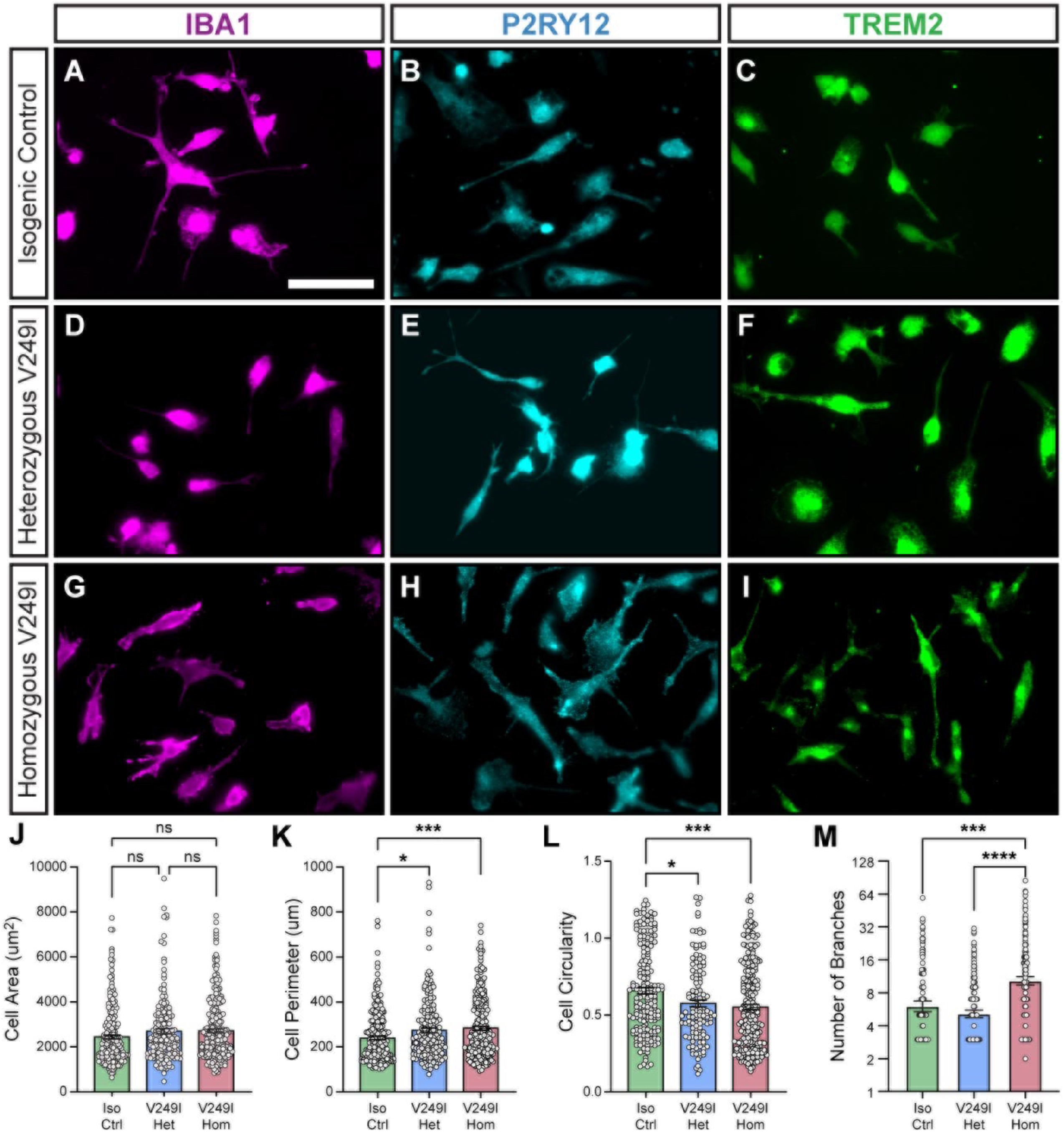
Characterization of hMGLs differentiated from human iPSCs. (AI) Wild type, heterozygous V249I, and homozygous V249I hMGLs all stained positive for characteristic microglial markers, including IBA1, P2RY12, and TREM2. (J) No differences were observed in cell areas across genotypes. Compared to controls, V249I heterozygous and homozygous hMGLs showed increased (K) perimeter (p=.0296, p=.0010) and (L) decreased circularity (p=.0219, p=.0002). (M) Additionally, homozygous V249I hMGLs demonstrated increased number of branches compared to both controls (p=.0001) and heterozygous V249I hMGLs (p<.0001). One-way ANOVA followed by Tukey’s multiple comparisons test. Scale bar: 50 µM.

### CX3CR1-V249I hMGLs demonstrate increased starvation-induced cell death and decreased proliferation

As fractalkine signaling plays a role in maintaining microglial survival (Boehme et al., 2000), and deficiency of fractalkine signaling has been shown to result in increased production of reactive oxygen species, leading to cell death (Arnoux and Audinat, 2015), we sought to assess whether the V249I SNP contributes to altered microglial survival. hMGLs were differentiated and then starved of supportive cytokines for varying lengths of time (Figure 3A-L), as previously described (McQuade *et al*., 2020), and cellular viability was determined with a Live/Dead cellular assay. While no differences in cell death were observed under basal conditions prior to cytokine starvation (Figure 3M), homozygous CX3CR1-V249I hMGLs demonstrated increased cell death compared to both heterozygous and isogenic controls after 24 hours (Figure 3N). After 48 hours of starvation, however, both heterozygous and homozygous CX3CR1-V249I hMGLs showed similarly increased levels of cell death compared to isogenic controls, suggesting a gene dosage-dependent effect.

**Figure 3.**
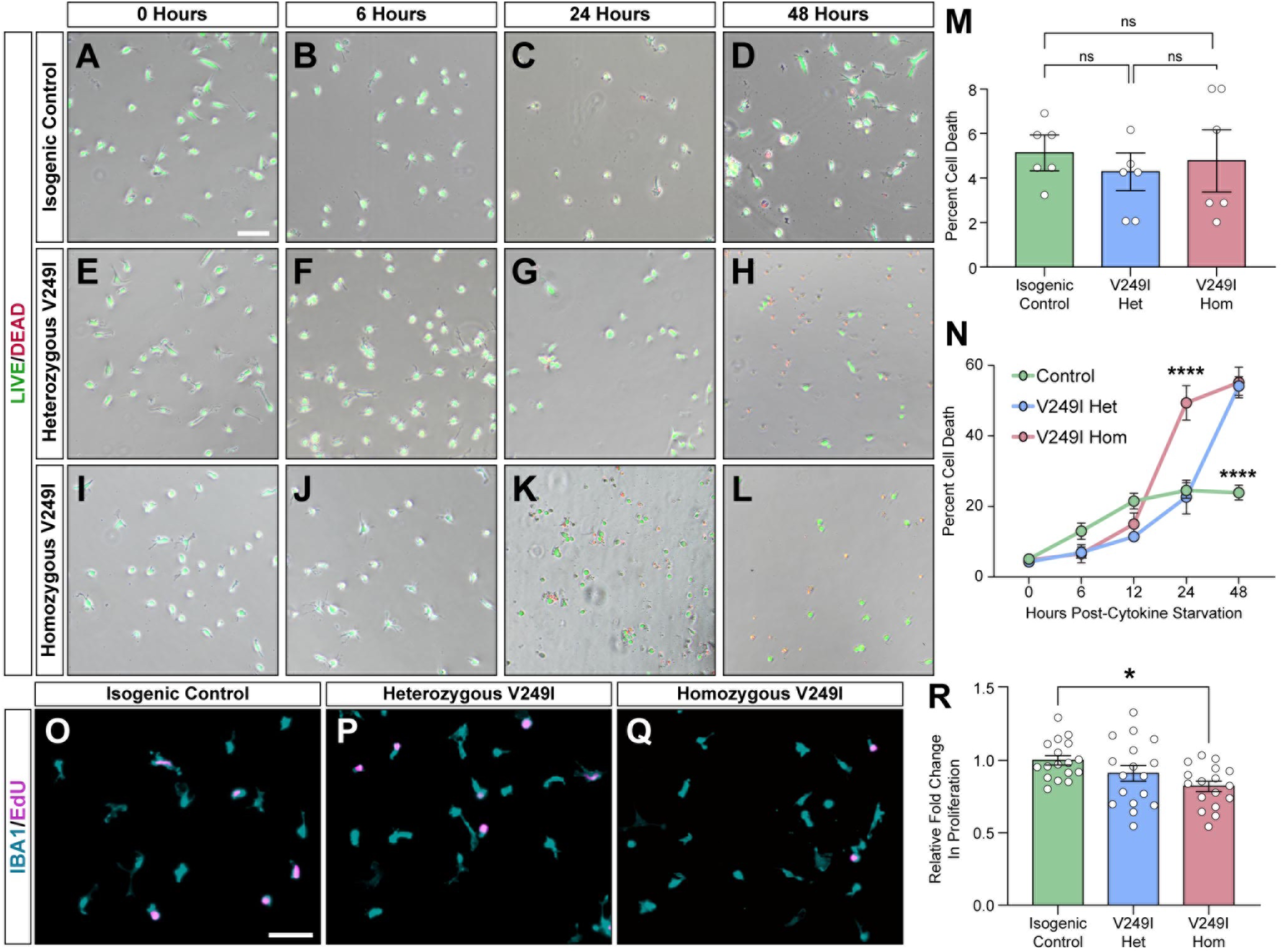
V249I hMGLs demonstrate increased susceptibility to starvation-induced cell death and decreased proliferation. (A-L) Control, heterozygous V249I, and homozygous V249I hMGLs were starved of supportive cytokines mCSF, TGFβ, and IL-34 for up to 24 hours. Live and dead cells were marked with the Life Technologies LIVE/DEAD™ Fixable Lime (506) Viability Kit. Scale bar: 200 µm. (M) At basal conditions, no differences were observed in percentage cell death based on genotype. (N) After 24 hours of starvation, homozygous V249I hMGLs showed increased percentage cell death compared to both controls (p<.0001) and heterozygous V249I hMGLs (p<.0001). After 48 hours of starvation, both heterozygous (p<.0001) and homozygous (p<.0001) V249I hMGLs demonstrated increased cell death compared to controls. (O-Q) At basal conditions, hMGLs were treated with the Life Technologies Click-It EdU kit for visualization of proliferative cells after 24 hours of Edu treatment. Scale bar: 250 µm. (R) Homozygous V249I hMGLs alone demonstrate decreased proliferation compared to controls (p=.0104) One-way (M and R) or two-way (N) ANOVA followed by Tukey’s multiple comparisons test.

Additionally, as fractalkine signaling leads to a more proliferative microglial state (Perros et al., 2007; White et al., 2010), we sought to determine whether the V249I SNP conferred alterations to microglial proliferation. Differentiated hMGLs were treated with 5-ethynyl 2’-deoxyuridine (EdU) for 24 hours to assess differences in the percentage of proliferative cells. We observed a decreased proliferation of hMGLs bearing the homozygous CX3CR1-V249I SNP compared to controls (Figure 3O-R), although no significant differences were observed in heterozygous CX3CR1-V249I hMGLs. To assess if V249I-related proliferation differences involved inflammatory pathways, cytokine secretion and gene expression were analyzed in hMGLs, revealing no genotype-dependent changes in pro- or anti-inflammatory cytokines (Supplemental Figure 3).

### CX3CR1-V249I hMGLs exhibit phagocytic deficits

An essential function of microglia is the uptake and degradation of toxic proteins (Colonna and Butovsky, 2017), and deficits in CX3CR1 signaling have been associated with dysfunctional phagocytosis (Bhaskar *et al*., 2010; Castro-Sanchez *et al*., 2019). To determine whether the CX3CR1-V249I variant affects microglial phagocytosis, hMGLs were treated with aggregated 488-tagged Aβ 1-42 (Figure 4A-C), and phagocytosis of this substrate was assessed via flow cytometry. Results demonstrated that while no significant differences were observed in the percentage of cells positive for phagocytosed Aβ at basal conditions (Figure 4D-G), both the heterozygous and homozygous V249I hMGLs demonstrated a decreased percentage of highly fluorescing cells (yellow highlighted area in Figure 4D-G, quantified in Figure 4L-M), suggesting a decreased, but not eliminated, capacity for V249I hMGLs to phagocytose Aβ. To further determine whether this effect was modulated by fractalkine (CX3CL1) signaling, cells were then pre-treated with fractalkine for 24 hours prior to Aβ exposure (Figure 4H-K). In hMGLs pre-treated with fractalkine, only homozygous V249I hMGLs demonstrated a decreased percentage of highly fluorescing cells, suggesting that the dysfunctional phagocytosis phenotype in heterozygous hMGLs was effectively rescued by fractalkine signaling (Figure 4N-O).

**Figure 4.**
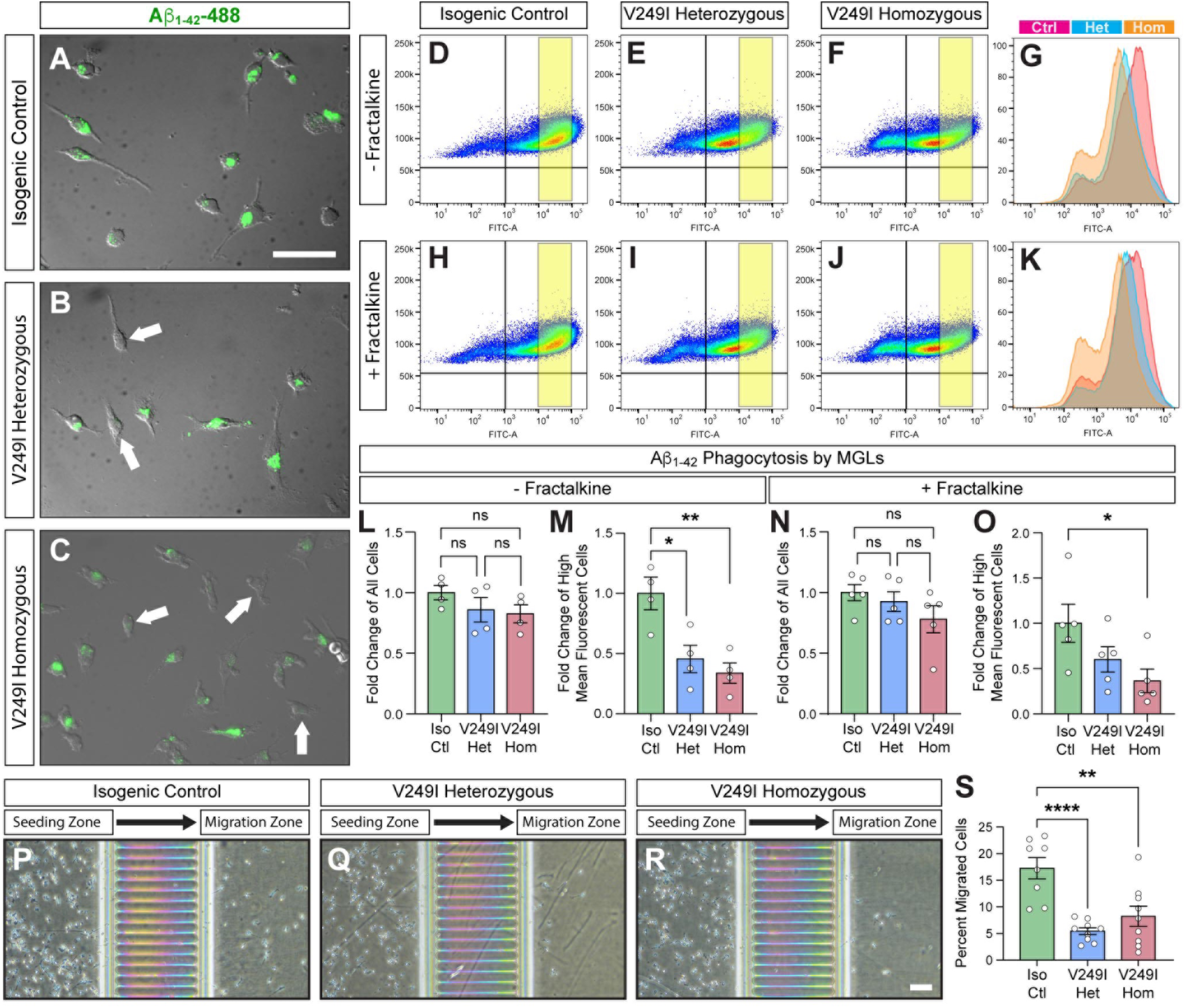
V249I hMGLs demonstrate decreased uptake of amyloid beta and migration. (A-C) hMGLs from all genotypes were treated with aggregated, 488-tagged Aβ 1-42 for one hour. (D-K) Flow sorting of cells with and without treatment of exogenous fractalkine 24 hours prior to Aβ addition. (L) At basal conditions, no differences were observed in percentage of Aβ+ cells. (M) Both heterozygous (p=.0199) and homozygous (p=.0066) V249I hMGLs had a lower percentage of highly fluorescing cells compared to controls. (N) With fractalkine treatment, no differences remained in the percentage of Aβ+ cells. (O) Homozygous V249I hMGLs alone demonstrate a decreased percentage of highly fluorescing cells compared to controls (p=.0442). (P-R) hMGLs were plated on one side of a microfluidic chamber and allowed to migrate across microfluidic grooves for 24 hours. (S) After 24 hours, both heterozygous (p=.0019) V249I hMGLs show a decreased percentage of migrated cells compared to controls. Scale bar equals 50 um in A and 100 um in R. One-way ANOVA followed by Tukey’s multiple comparisons test.

### CX3CR1-V249I hMGLs exhibit migratory deficits

Fractalkine signaling is additionally responsible for modulating microglial migration and environmental surveillance (Poniatowski et al., 2017; Sheridan and Murphy, 2013). Previous work has demonstrated that decreased CX3CR1 function has led to deficient migration (Castro-Sanchez *et al*., 2019; Maciejewski-Lenoir et al., 1999), and dysfunctional migration may partially explain reduced uptake of Aβ observed in hMGLs with the V249I genotype. Thus, to identify deficits in microglial migration based on the presence of the V249I SNP, we seeded hMGLs into a microfluidic platform to assess microglial migration through microfluidic channels into the otherwise empty contralateral side (Figure 4P-R). After 24 hours of spontaneous migration, both heterozygous and homozygous V249I hMGLs showed a decreased migratory capacity into the opposing chamber compared to wild type controls (Figure 4S), suggesting a dysfunction in microglial motility with the V249I SNP.

### Transcriptional profiling of CX3CR1-V249I hMGLs

The CX3CR1-V249I SNP may confer dysfunctional microglial phenotypes through various mechanisms. Thus, to further explore how this gene variant alters hMGLs more globally, we explored their transcriptional state when derived from isogenic control cells as well as from the heterozygous or homozygous genotype via RNA-seq (Figure 5). A comparison between heterozygous CX3CR1-V249I and isogenic control hMGLs revealed several downregulated genes linked to cell adhesion (PCDHGB4), migration (S100A10) and cell cycle (Gas8, DNAJA4, TSPYL5, HPDL) in the heterozygous CX3CR1-V249I hMGLs. Interestingly, most of the upregulated genes found in this variant were related to inflammatory pathways (such as IRAK4, ADGRG1 and PRKCH) (Figure 5A-C), suggesting a deregulated and inflammatory potential in heterozygous cells. When we compared the transcriptional profile of homozygous CX3CR1-V249I with isogenic control hMGLs, most of the differentially expressed genes were similar to the previous comparison, especially those upregulated genes related to inflammatory signaling (Figure 5D-F). Despite the similarities identified between heterozygous and homozygous hMGLs compared to control samples, we next wanted to determine differentially expressed genes that uniquely characterize each variant genotype. While similarities in phenotypes arose between the V249I heterozygous hMGLs and V249I homozygous hMGLs, there were distinct differences observed in this study, with homozygous hMGLs appearing to have more severely affected phenotypes. When comparing heterozygous and homozygous V249I hMGLs, we observed that genes related to immune response were most upregulated in both genotypes, but with distinct signatures, as the homozygous cells exhibit upregulation of GPR39, CCL22, SOCS1, while heterozygous cells show higher levels of SLC15A4 (Figure 5G-I). These results suggest some level of dysfunctional immune modulation between these lines.

**Figure 5:**
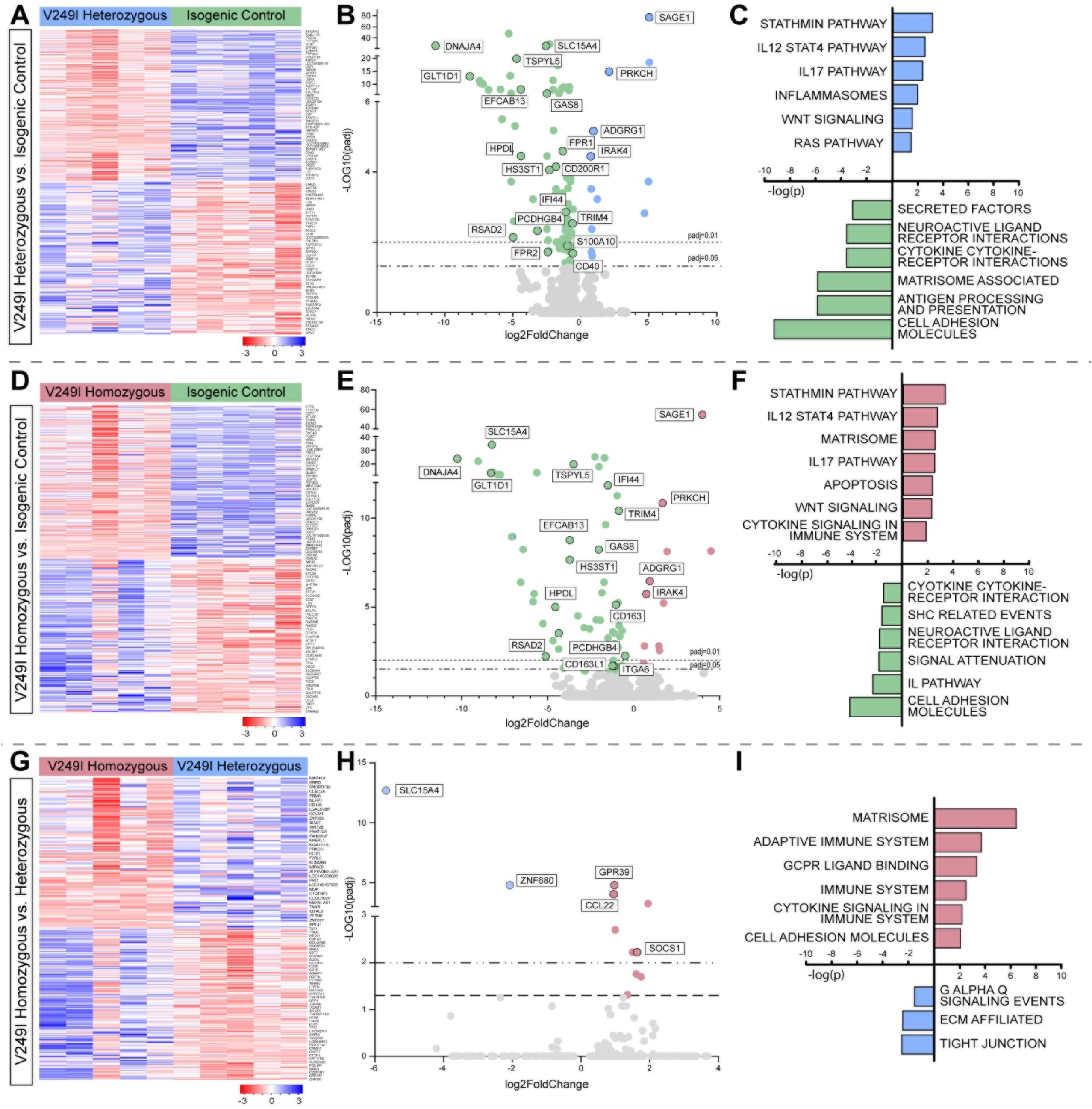
RNA sequencing reveals transcriptional changes in V249I hMGLs. Heat map demonstrates differential gene expression patterns in comparisons between (A) heterozygous V249I hMGLs vs control, (D) homozygous V249I hMGLs vs control, and (G) homozygous V249I hMGLs vs heterozygous. Each column represents one sample, and each row represents one gene. The up-and down-regulated genes are blue and red colored, respectively. (B, E, H) Volcano plot exhibiting differentially expressed genes across comparisons. (C, F, I) GO and pathway analysis based on differential gene expression revealed a number of altered pathways across comparisons.

Additionally, pathway enrichment analyses identified numerous cellular pathways differentially modulated due to the V249I genotype, including an upregulation of genes linked to the stathmin pathway, IL12 STAT4 pathway, and IL17 pathway. Moreover, most of the downregulated genes in V249I variants were related to neuroactive ligand receptor interaction, cytokine-cytokine interaction, and cell adhesion molecules (Figure 5C-F). Interestingly, while V249I heterozygous hMGLs demonstrated downregulation of interferon gamma signaling pathways, GCPR ligand binding, and matrisome pathways, these pathways were upregulated in V249I homozygous V249I hMGLs compared to controls (Figure 5I), suggesting a specific transcriptional signature depending on the allele that may explain the phenotypic differences observed in this study.

### Morphological changes upon co-culture of iPSC-derived microglia and neurons

While results described above have demonstrated that the CX3CR1-V249I SNP has cell autonomous effects on hMGL phenotypes, previous studies have also demonstrated that hMGLs adopt more physiologically relevant phenotypes when grown in an environment more similar to those within the brain, in which non-cell autonomous effects can also be observed (Popova et al., 2021). To assess changes to hMGL phenotype based upon CX3CR1 genotype in a co-culture system, hMGLs were cultured with iPSC-derived neurons, the primary producers of the CX3CL1 ligand, derived by induced NGN2 expression (Figure 6A), (Hatori et al., 2002). Co-culture with neurons increased hMGLs complexity in all genotypes compared to hMGLs grown alone (Figure 6B-E), with increased maximum branch length for all genotypes in co-culture with neurons compared to hMGLs grown alone. Across genotypes, we also observed that in the presence of neurons, hMGL homozygous for the V249I variant exhibited significantly more branches per cell as well as more junctions per cell (Figure 6F-G). To determine if hMGLs with or without the V249I genotype conversely modulated neuronal phenotypes, we then assessed neuronal morphologies when grown in co-culture with hMGLs. We observed that iPSC-derived neurons co-cultured with heterozygous V249I hMGLs displayed an increased number of neurites compared to the effect promoted by control hMGLs, while homozygous hMGLs failed to modulate a similar increase in neuronal complexity (Figure 6H-K).

**Figure 6:**
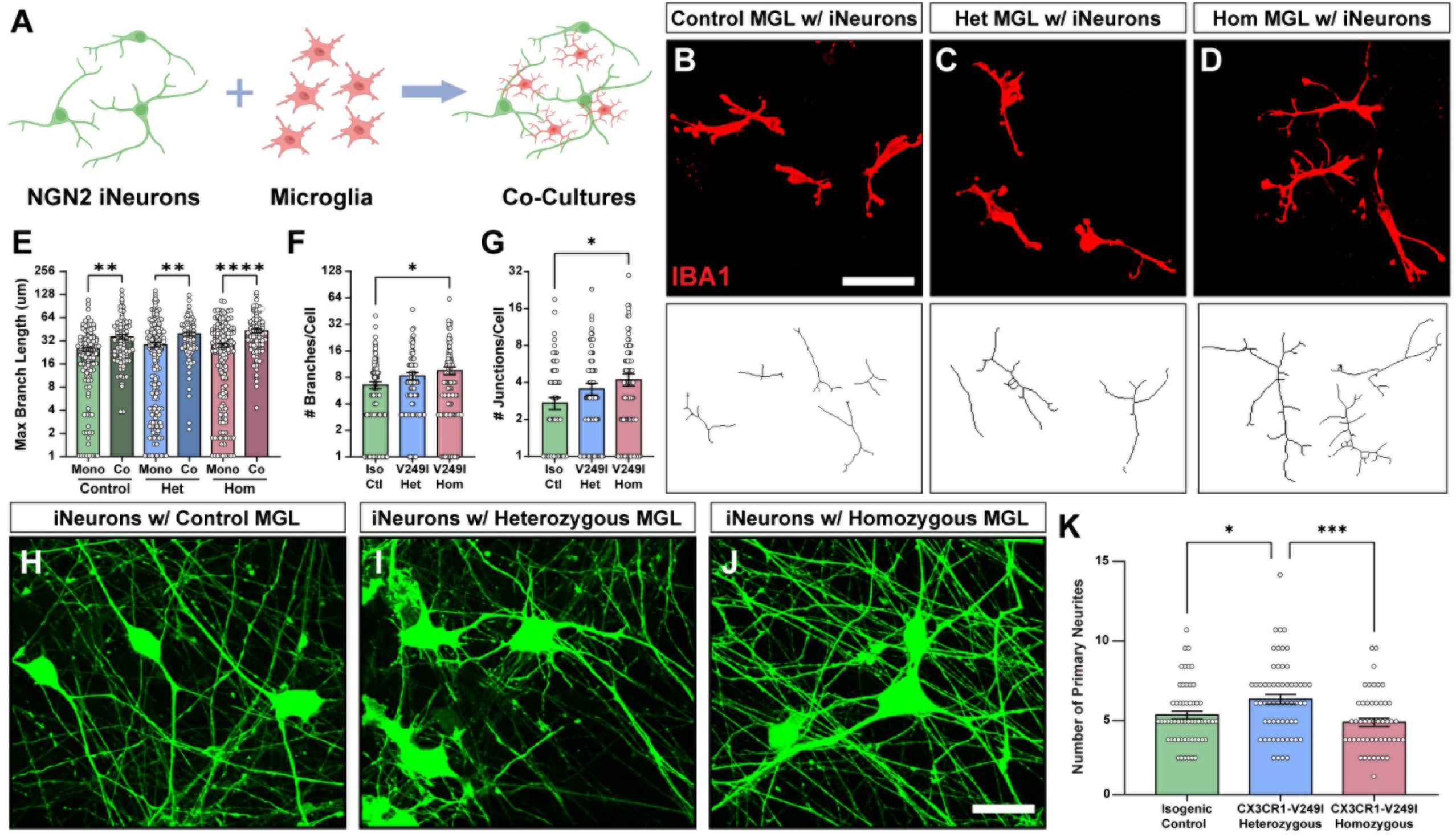
V249I SNP modulates cell autonomous and non-cell autonomous effects in co-culture systems. (A) Schematic of co-culture strategy of cortical neurons and hMGLs. (B-D) Representation of co-cultured hMGL morphology after staining with IBA1 across all genotypes, as well as representation of hMGLs skeletonization per condition. Scale bar: 50 µm (E) Compared to monoculture conditions, co-culture with neurons promoted increased maximum branch length for control (p=.001), heterozygous (p=.0045), and homozygous (p<.0001) V249I hMGLs. (F-G) In co-culture conditions, homozygous V249I hMGLs demonstrated an increased number of branches (p=.0204) and junctions (p=.0182) compared to controls. (H-J) Representative images of eGFP-expressing neurons co-cultured with hMGLs from control, heterozygous V249I, and homozygous V249I genotypes. Scale bar: 20 µm. (K) Compared to co-culture with both control (p=.0250) and homozygous (p=.0009) V249I hMGLs, neurons co-cultured with heterozygous V249I hMGLs demonstrated an increased number of primary neurites. One-way ANOVA followed by Tukey’s multiple comparisons test.

We then sought to determine if MGLs could modulate the activity of neuronal populations, either with or without the CX3CR1-V249I genotype. To examine this, we developed cortical organoids from iPSCs and then seeded hMGLs into these organoids (Figure 7A-E). Organoids seeded with hMGLs were then plated onto MEA plates, and electrical activity of neurons within the organoids was analyzed (Figure 7F-L). Results demonstrated that heterozygous V249I hMGLs were able to modulate neuronal activity by increasing the number of spikes, mean firing rate, and number of bursts. These results suggest that hMGLs carrying the heterozygous CX3CR2-V249I variant modulated morphological and functional changes to neuronal populations, potentially contributing to neurodegenerative phenotypes.

**Figure 7:**
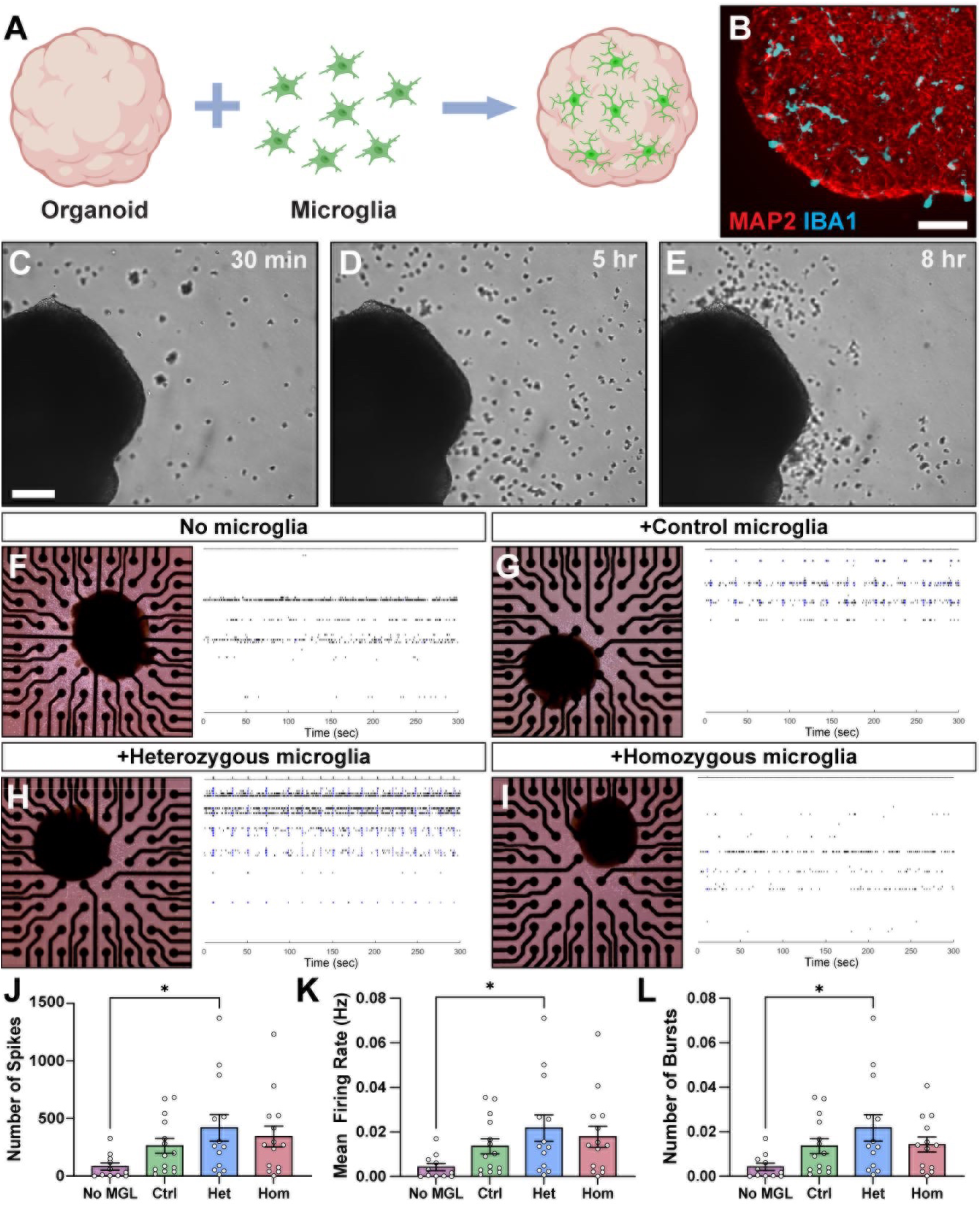
V249I hMGLs promote functional alterations of neurons in cortical organoids. (A-B) Schematic of hMGLs seeded into cortical organoids, as well as representative image of MAP2 and IBA-1 demonstrating the presence of neurons and microglia, respectively, in the organoid. Scale bar: 200 µm. (C-E) Representative time lapse images of hMGLs migrating into cortical organoids after 30 min, 5 hours, and 8 hours. Scale bar: 200 µm. (F-I) Representation of cortical organoids containing hMGLs attached to microelectrode plates, as well as representation of raster plots from organoids containing each genotype of microglia. Black bars represent spikes and blue bars represent bursts. (J-L) Compared to organoids not containing hMGLs, only heterozygous hMGLs promoted increased number of spikes (p=.0456), mean firing rate (p=.0452), and number of busts (p=.0231) in organoids. One-way ANOVA followed by Tukey’s multiple comparisons test.

## DISCUSSION

The development of model systems capable of identifying physiologically relevant phenotypes in human cells is critical for the development of therapeutic interventions for neurodegenerative diseases. The fractalkine axis regulates key microglial functions, including migration, cytokine release, neuroprotection and cell survival, and phagocytosis (Arnoux and Audinat, 2015; Boehme *et al*., 2000; Lastres-Becker *et al*., 2014). Additionally, impaired fractalkine signaling exacerbates Alzheimer’s Disease pathology, with reduced function of the fractalkine receptor linked to worsened Tau pathology and cognitive decline (Bhaskar *et al*., 2010; Castro-Sanchez *et al*., 2019; Puntambekar *et al*., 2022).

Previous work has suggested that the CX3CR1 receptor plays an important role in regulating microglia survival and response to stress (Boehme *et al*., 2000; Castro-Sanchez *et al*., 2019). To investigate whether the V249I polymorphism affects these functions, hMGLs were starved of supportive cytokines (mCSF, TGFβ, IL-34), mimicking the loss of neuronal support in late-stage neurodegeneration. Interestingly, while no differences in cell death were observed at basal conditions, cytokine starvation resulted in early and pronounced cell death in homozygous V249I hMGLs, with heterozygous hMGLs exhibiting similar cell death at later time points. These results suggest the V249I SNP reduces microglial resilience to cytokine loss. Similar vulnerability has been reported in TREM2-deficient hMGLs (McQuade *et al*., 2020), emphasizing the important role of microglial-environment interactions in supporting survival and potentially contributing to neurodegenerative-phenotypes and pathology.

Studies on microglia lacking CX3CR1 function have shown mixed effects on phagocytosis (Bhaskar *et al*., 2010; Castro-Sanchez *et al*., 2019; Lee and Landreth, 2010). In this work, we observed that the V249I SNP did not affect amyloid beta uptake initiation but reduced overall phagocytic capacity. Interestingly, this effect can be overcome by treatment with exogenous ligand in the heterozygous, but not the homozygous, hMGLs. As previous data has suggested that heterozygous hMGLs do not confer risk in the development of AD (Lopez-Lopez *et al*., 2018), the ability of fractalkine to modulate an effect on Aβ phagocytosis may explain why heterozygous hMGLs remain fully functional. Our findings support the idea that heterozygous V249I hMGLs retain effective amyloid clearance in physiological conditions, while homozygous cells show impaired uptake.

To our knowledge, this is the first study to examine transcriptional changes due to the V249I polymorphism in human microglia in the absence of the influence of other cell types, allowing a direct analysis of cell autonomous changes due to this variant. Many of the differentially expressed genes identified in this work were related to immune cell function, with both pro-inflammatory and anti-inflammatory genes being highlighted in our data. Notably, the stathmin pathway, a microtubule regulating protein that has been implicated in the modulation of microglial inflammatory responses (Bsibsi et al., 2010), was downregulated in both V249I genotypes. We also observed alterations in IL-12/STAT4 signaling, a pro-inflammatory pathway active in human microglia (Taoufik et al., 2001). Of particular note, IRAK4, a key mediator of toll-like receptor signaling and inflammation, was upregulated in both heterozygous and homozygous V249I hMGLs. Its increased activity has been linked to impaired amyloid beta clearance and AD pathology (Bai et al., 2023; Cameron et al., 2012). Additionally, an upstream signaling molecule in this pathway, SLC15A4, was observed downregulated in both heterozygous and homozygous hMGLs, with a downregulation even more pronounced in heterozygous hMGLs. SOCS1, an inhibitor of this pathway (Pereira and Gazzinelli, 2023), was elevated in in the homozygous compared to heterozygous hMGLs. These expression patterns suggest altered TLR9 signaling, which may be implicated in some of the observed AD-related phenotypes, with differential modulation separating heterozygous and homozygous V249I hMGLs.

As microglial pathology is a keystone feature of many neurodegenerative diseases including Alzheimer’s Disease, its impact on neuronal health remains critical for understanding disease progression and therapeutic potential. To investigate the effects of hMGLs on neurons, control and V249I hMGLs were co-cultured with both 2D neurons and 3D cortical organoids. Heterozygous V249I hMGLs uniquely promoted neurite outgrowth and neuronal hyperexcitability, possibly due to CX3CR1-related calcium signaling changes (Deiva et al., 2004). Further, our RNA-seq analysis revealed several genes involved in calcium signaling and neurotransmitter modulation. Both heterozygous and homozygous hMGLs exhibited reduced expression of Glt1d1, a critical gene for glutamate reuptake that is dysregulated in Alzheimer’s Disease (Srinivasan et al., 2020) and whose downregulation may contribute to neuronal hyperexcitability. Additionally, fractalkine receptor signaling is known to delay maturation of glutamate receptors (Hoshiko et al., 2012), making the lack of neuronal hyperexcitability in homozygous V249I hMGLs co-cultures unexpected. Notably, heterozygous V249I hMGLs alone exhibited increased expression of KIF17, an NMDA transporter (Liu et al., 2022), and promoted neurite outgrowth in co-culture with cortical neurons. Given that receptor loss reduces neurotrophic gene expression, homozygous hMGLs may exert neurotoxic effects that mask excitability changes. Future work should clarify how this SNP impacts calcium signaling and neuronal health.

Homozygous V249I hMGLs showed decreased proliferation at maturity compared to controls, aligning with prior studies linking fractalkine signaling to microglial proliferation (Perros *et al*., 2007; Tang et al., 2015; White *et al*., 2010). Although pro-inflammatory microglia typically exhibit increased proliferation (Orihuela et al., 2016), this was not observed, likely due to the cell-autonomous nature of the experiments. Previous studies have shown that *in vivo*, microglial cytokine release activates astrocytes creating a feedback loop that amplifies inflammation (Liddelow et al., 2017), which fractalkine signaling normally helps resolve (Gonzalez-Prieto et al., 2021). It is possible that this cell non-autonomous feedback is essential for uncovering inflammatory profile changes in hMGLs bearing the V249I SNP. The absence of this intercellular cross-talk may explain the lack of inflammatory phenotypes observed in our study, highlighting the need for future studies involving multicellular systems to assess inflammatory modulation by the V249I genetic variant.

The utilization of iPSC models provides a unique and essential opportunity to uncover phenotypes resulting from genetic variants in human cells. These platforms enable the study of cell type specific functional changes without the confounding influence of other cell types. Additionally, co-culture models further extend this by revealing how genetic variants affect intracellular interactions. However, challenges remain in developing physiologically relevant iPSC models, including incorporating diverse cell types, addressing aspects of aging, and accounting for genetic variability across individuals. While this study uncovers several important roles of the CX3CR1-V249I SNP in microglial dysfunction, it cannot completely capture the complexity of the *in vivo* environment, including the influence of other cell types of the brain, spatiotemporal patterning, and vascularization. Nonetheless, this study provides a critical foundation for future research using more advanced and complex systems, which may include 3D co-culture iPSC-derived platforms, the incorporation of AD-gene bearing organoids, or implantation into murine models, to further elucidate the impact of the CX3CR1-V249I SNP.

## METHODS

### Maintenance and expansion of iPSCs

The WTC11 (Coriell GM25256) was used for all studies. These cells were karyotyped and regularly screened for mycoplasma using a commercially available kit (Millipore-Sigma). iPSCs were maintained in 6-well plates on a Matrigel substrate and media was changed every two days with mTeSR+ (StemCell Technologies). Colonies were grown to approximately 60-70% confluency and passaged using ReLeSR (StemCell Technologies) and replated at a 1:20 dilution or used for further differentiation to hMGLs or for organoid differentiation.

### CRISPR/Cas9 Editing of the V249I Gene Variant

CRISPR/Cas9 gene editing was accomplished utilizing previously described techniques (VanderWall et al., 2020) to introduce the CX3CR1-V249I SNP into these cell lines. Briefly, a gRNA sequence was designed targeting the V249I position and was synthesized along with an HDR template (Genscript). These plasmids were electroporated alongside a Cas9-GFP plasmid into iPSCs using the Neon Transfection System (Life Technologies). Electroporated iPSCs were seeded onto Matrigel-coated 6-well plates with ROCK Inhibitor. Two days after electroporation, fluorescence-activated cell sorting (FACS) was used to select transiently GFP-positive cells containing the Cas9-GFP plasmid to isolate those cells that had been properly electroporated. After an additional passage, iPSCs were plated at low density to allow for clonal expansion. Isolated clonal populations were screened for the anticipated removal of the AclI restriction enzyme site and then further Sanger sequenced to confirm genotype. Additionally, the top 5 most likely off target editing sites were determined using off-target screening tools (Synthego), and each of these regions were Sanger sequenced and confirmed to be identical to the isogenic control (primer sequences found in Supplemental Table 2).

### Differentiation of hMGLs

Differentiation of iPSCs into human microglia-like cells (hMGLs) was conducted using previously established protocols (Abud et al., 2017; McQuade et al., 2018). Briefly, iPSC colonies were grown on Matrigel to approximately 70% confluency, at which point cells were detached from the plate using ReLeSR reagent and small cellular aggregates were plated onto a fresh 6-well plate coated with Matrigel at a density of ∼30 colonies per well at a size of 150-200 microns per colony. The STEMdiff Hematopoietic Kit (StemCell Technologies) was used to differentiate these cells into hematopoietic progenitor cells (HPCs). On Day 12 of differentiation, HPCs were collected from the media and frozen in Bambanker Freezing Medium (Fujifilm Irvine Scientific) at a density of 500,000 cells/1mL and cryopreserved in liquid nitrogen for subsequent differentiation into hMGLs. To differentiate HPCs into MGLs, control, heterozygous CX3CR1-V249I, and homozygous CX3CR1-V249I HPCs were thawed in parallel from liquid nitrogen into two wells of a 6 well plate for each cell line. Microglia Basal Media was used for differentiation, which consisted of: DMEM/F12, 2% Insulin-Transferrin-Selenium, 2% B27, 1% Anti Anti, 1% MEM NEAA, 1% Glutamax, 0.5% N2, 0.2mg/mL insulin, and 400uM monothioglycerol, and further supplemented with 100ng/mL IL-34, 50ng/mL TGF-β1, and 25ng/mL mCSF. HPCs were transitioned to this new media upon thaw and fed every 48 hours. After 25-28 days of maturation, hMGLs were collected for assays.

### Immunostaining

hMGLs were collected and plated on coverslips coated with Poly-D-Lysine and Vitronectin at a density of 30,000 cells per 12 mm coverslip. Cells were fixed in 4% paraformaldehyde for 15 minutes and washed three times in PBS. hMGLs were then permeabilized in 0.2% Triton X-100 for 10 minutes and blocked in 5% bovine serum albumin (BSA) in PBS for 20 minutes. Subsequently, hMGLs were incubated in primary antibody in 5% BSA for 30-60 minutes at room temperature. After primary antibody incubation, cells were washed three times in PBS, and then incubated with secondary antibody in 5% BSA for 30-60 minutes. After three final washes with PBS, coverslips were mounted onto microscope slides and imaged using a Leica DM5500 fluorescence microscope. Antibodies used for these studies can be found in Supplemental Table 1.

### Cell Viability Assays

Following hMGL differentiation, cells were harvested and replated onto 4 wells of a 96-well plate coated with Vitronectin (Life Technologies). After 24 hours to allow adherence, a full media change was performed into fresh medium lacking IL-34, TGF-β, and mCSF for 6, 12, 24, or 48 hours. The Live/Dead Cell Imaging Kit (Life Technologies) was applied following manufacturer’s instructions, and cells were imaged at 10x using a Nikon Eclipse TS2R-FL microscope with a Nikon DS-Ri2 camera. The percentage of cell death was calculated as the number of dead cells (red) as a ratio to the total number of cells in each image. A minimum of three images were taken per biological replicate across four separate differentiations of cells (biological replicates).

### EdU Proliferation Assay

hMGLs were harvested and replated onto 4 wells of a 96-well plate coated with Vitronectin (Life Technologies). After 24 hours to allow adherence, hMGLs were treated with EdU from the Click-It EdU Kit (Life Technologies C10637) for 24 hours. Cells were then fixed with 4% paraformaldehyde, the rest of the assay was performed according to manufacturer’s instructions, and cells were counterstained with IBA1 to confirm microglial phenotype as well as a DAPI nuclear counterstain. Each well was imaged at 4x using a Nikon Eclipse TS2R-FL microscope with a Nikon DS-Ri2 camera. The percentage of proliferating cells was quantified as the number of EdU-positive nuclei compared to the total number of DAPI-positive nuclei in each image. A minimum of three images were taken per biological replicate across four separate differentiations of cells (biological replicates).

### Phagocytosis of Amyloid Beta and Flow Cytometric Analysis

To assess the phagocytic capability of hMGLs across genotypes, a 488-tagged amyloid beta 1-42 (Anaspec) was initially reconstituted according to manufacturer instructions and one day prior to treatment, Aβ was mixed 1:1 with sterile MilliQ water and pipetted vigorously. It was then incubated overnight at 37 degrees Celsius for aggregation as previously described (Kleinberger et al., 2014; Lin et al., 2018; Xiang et al., 2016), and cells were then treated with the aggregated amyloid beta 1-42 at a final concentration of 2ug/mL and incubated at 37 degrees C for 1 hour prior to analysis by flow cytometry. Phagocytosis of aggregated Aβ1-42-488 by iPSC microglia was assessed by flow cytometry as described previously (Puntambekar *et al*., 2022). Briefly, 1x10^6^ iPSC microglia were incubated with human Fc-Block (BD Pharmigen, Cat # 564219) to eliminate non-specific antibody binding. Cells were stained using DRAQ7 (Biolegend, Cat # 424001) for exclusion of dead cells, and CD11b-PE (Biolegend, Cat # 101208, Clone M1/70) for 30 mins on ice. All experiments were performed using appropriate, bead-based single-colored controls to correct for non-specific bleed through due to spectral overlap. All analysis of flow-cytometry data was done using the Flow Jo analysis software.

### Microglial migration assay

hMGLs were collected from one well of a 6 well plate and replated onto the left chamber of the Xona Microfluidic chips in 10uL of media. After 24 hours of incubation, microfluidic devices were fixed in 4% PFA for 15 minutes and washed with PBS. Three areas of each microfluidic were imaged, and the number of microglia on each side of the chamber was quantified. The percentage of migrated cells was calculated as the number of cells in the right chamber divided by the total number of microglia in each image. Three images were taken per microfluidic across three separate differentiations of cells (biological replicates).

### RNA Extraction and RNAseq

Differentiated MGLs were collected and centrifuged, and then lysed using extraction buffer from the PicoPure RNA Extraction Kit. The PicoPure Kit (Life Technologies) was used to isolate total RNA following manufacturer’s instructions. Collected RNA was analyzed for quality and concentration using the Agilent Bioanalyser 2100 at the Indiana University Center for Medical Genomics. Approximately 30 million reads per library were generated. The sequencing data were next assessed using FastQC (Wingett and Andrews, 2018) and then mapped to the human genome (GRCH38) using STAR RNA sequencing aligner (Dobin et al., 2013). Differentially expressed genes were tested by using DESeq2 with a false discovery rate < 0.05 as the significant cutoff (Love et al., 2014). Pathway enrichment analysis were conducted by hypergeometric test against human gene ontology and MsigDB v.6 canonical pathways, with p < 0.01 as the significant cutoff (Subramanian et al., 2005). The RNA sequencing experiments reported in this paper have been deposited in the Gene Expression Omnibus database, www.ncbi.nih.gov/geo (accession no. GSEXXXXX).

### NGN2-based Neuronal Differentiation and Co-Culture with Microglia

iPSC-derived neurons were differentiated via delivery of NGN2, as previously described (Zhang and Zhang, 2021). Briefly, plasmids were obtained from Addgene, including pLV-TetO-hNGN2-eGFP-Puro (item #79823) and FUdeltaGW-rtTA (item #19780). Lentiviral particles were generated using these plasmids, which were then used to transduce iPSCs to generate iNeurons, as previously described (Ho et al., 2016). Neuronal differentiation was then induced by doxycycline treatment (10mg/mL), and neuronal cells were selected with puromycin (10mg/mL) the next day. iNeurons were then harvested on day 4 following doxycycline treatment and either frozen or re-plated for experimental use on dishes coated with poly-ornithine (20 ug/ml) and laminin (5 ug/ml), followed by coating with Matrigel (42 ug/cm^2^) at a density of 50,000 cells per well of a 24 well plate. Cells were maintained in complete Neurobasal Media supplemented with 2% B27 Supplement, ROCK Inhibitor, 10mg/mL Puromycin, and 10mg/mL Doxycycline Hyclate for three days. After 3 days, hMGLs were added to these cultures at a density of 10,000 cells per well, and media was transitioned to complete BrainPhys with IL-34, TGF-β, and mCSF. Cells were fed every 48 hours for one week of co-culture, at which time coverslips were fixed in 4% paraformaldehyde and stained for IBA1 or MAP2.

### Differentiation of iPSC-derived cortical organoids and seeding with microglia

Differentiation of iPSCs into cortical organoids was conducted using previously established protocols (Fligor et al., 2020; Harkin et al., 2024; Li and Zhang, 2006) with slight adjustments. Briefly, iPSC colonies were grown on Matrigel to approximately 80% confluency and lifted using Dispase (2 mg/ml) at 37 degrees for 25 minutes. Colonies were transferred to media containing mTeSR+ and Neural Induction Media (NIM), consisting of DMEM/F12, 1% MEM NEAA, 1% Anti-Anti, 1% N2, and heparin (2 ug/ml). On day 6 of differentiation, embryoid bodies (EBs) were treated with 200uM of LDN-193189 to direct differentiation to a cortical lineage. On day 8, EBs were plated into 6 well plates containing 25% fetal bovine serum for 24 hours and maintained in NIM with media changes every 2-3 days until day 16. At this point, colonies were manually lifted from 6 well plates and transferred to Neural Maturation Media (NMM), consisting of DMEM/F12 (3:1), 2% B27, 1% MEM nonessential amino acids, and 1% Anti-Anti, and grown in suspension in 100mm petri dishes. NMM was changed every 2-3 days until organoids reached between day 40-50 of differentiation for assays. At this point, organoids were plated into a 96-well U-Bottom Low Adhesion Plate. hMGLs were added to each well of the 96 well plate containing a single sphere at a density of 10,000 MGLs/well and transitioned to complete BrainPhys media, consisting of BrainPhys Neuronal Media, 2% B27 Supplement, 1% N2, 1% Glutamax, 1% Anti-Anti. Cultures were supplemented with 20ng/mL BDNF, and 20ng/mL GDNF, 100ng/mL IL-34, 50ng/mL TGF-β1, and 25ng/mL mCSF for 3 days prior to further analyses.

### Electrophysiological recordings via multielectrode array

3 days following the seeding of microglia into cortical organoids, spheres were plated on to a laminin-coated 6-well Axion Biosystems MEA plate for one week with media changes every two days. After one week of culture on the MEA plate, organoid activity was analyzed using the Axion Biosystems Maestro Edge system. As organoids did not necessarily cover all electrodes on the plate, data points were normalized to the total number of covered electrodes. One to three organoids per biological replicate were included in analysis across six separate differentiations of cells (biological replicates).

### Morphological Measurements and Statistical Analyses

Images of microglia and/or neurons were acquired on either a Leica DM5500 fluorescence microscope or a Nikon Eclipse TS2R-FL microscope with a Nikon DS-Ri2 camera. Individual cells were traced using ImageJ with analysis of traced paths for area and perimeter. Circularity was taken as 4*pi(area/perimeter^2^). At least 39 cells per condition across three separate differentiations of cells (biological replicates) were quantified. For measurements of microglial morphology, the ImageJ AnalyzeSkeleton plug in was used with adjustment of threshold to exclude background and binary processing of skeletonization. At least 60 cells per condition across three separate differentiations of cells (biological replicates) were quantified. Data in all experiments was represented as mean ± SEM and n represents the number of technical replicates across all experiments. Statistical comparisons were conducted by either Student’s t-test or ANOVA with Tukey post hoc test using GraphPad Prism 9. Statistically significant differences were defined as *p*<0.05 in all experiments. All experiments included at least three independent differentiations of cells (biological replicates).

## Supporting information

Microglia Organoid

## ACKNOWLEDGMENTS

We thank Tracy Young-Pearse and Christina Muratore for assistance with NGN2-induced neuronal cultures. Grant support was provided by the National Institute for Aging (RF1AG069425 to BTL and JSM). Support for this project was also provided by a supplement award (RF1AG069425-01S1). KDT was also supported by an institutional training grant (T32AG071444) as well as a fellowship from the National Institute for Aging (F30AG084304). Schematics used in some figures created with BioRender.

## Contributions

KDT, CG, SSP, SJB, BTL, and JSM designed the research. KDT, JH, LDV, MH, YF, PW, LMB performed the research. KDT, MH, YF, TYL, CG, CZ, SSP, SJB, BTL, JSM analyzed data. KDT, CG, and JSM wrote the paper.

## Availability of data and materials

Data pertaining to this study that is not directly available within the article will be made available from the corresponding author upon reasonable request. Transcriptional data will be uploaded as a GEO Dataset.

## Declaration of interests

The authors declare no competing interests.

## SUPPLEMENTAL INFORMATION

**Supplemental Table 1:** Primary Antibodies used in this study.

| Primary Antibody | Species | Dilution | Company | Catalog Number |
| --- | --- | --- | --- | --- |
| IBA1 | Rabbit | 1:500 | Fujifilm Irvine Scientific | 019-19741 |
| P2RY12 | Rabbit | 1:200 | Sigma Aldrich | HPA014518 |
| TREM-2 | Goat | 1:100 | R&D Systems | AF1828 |
| MAP2 | Mouse | 1:200 | Synaptic Systems | 188011 |

**Supplemental Table 2:**
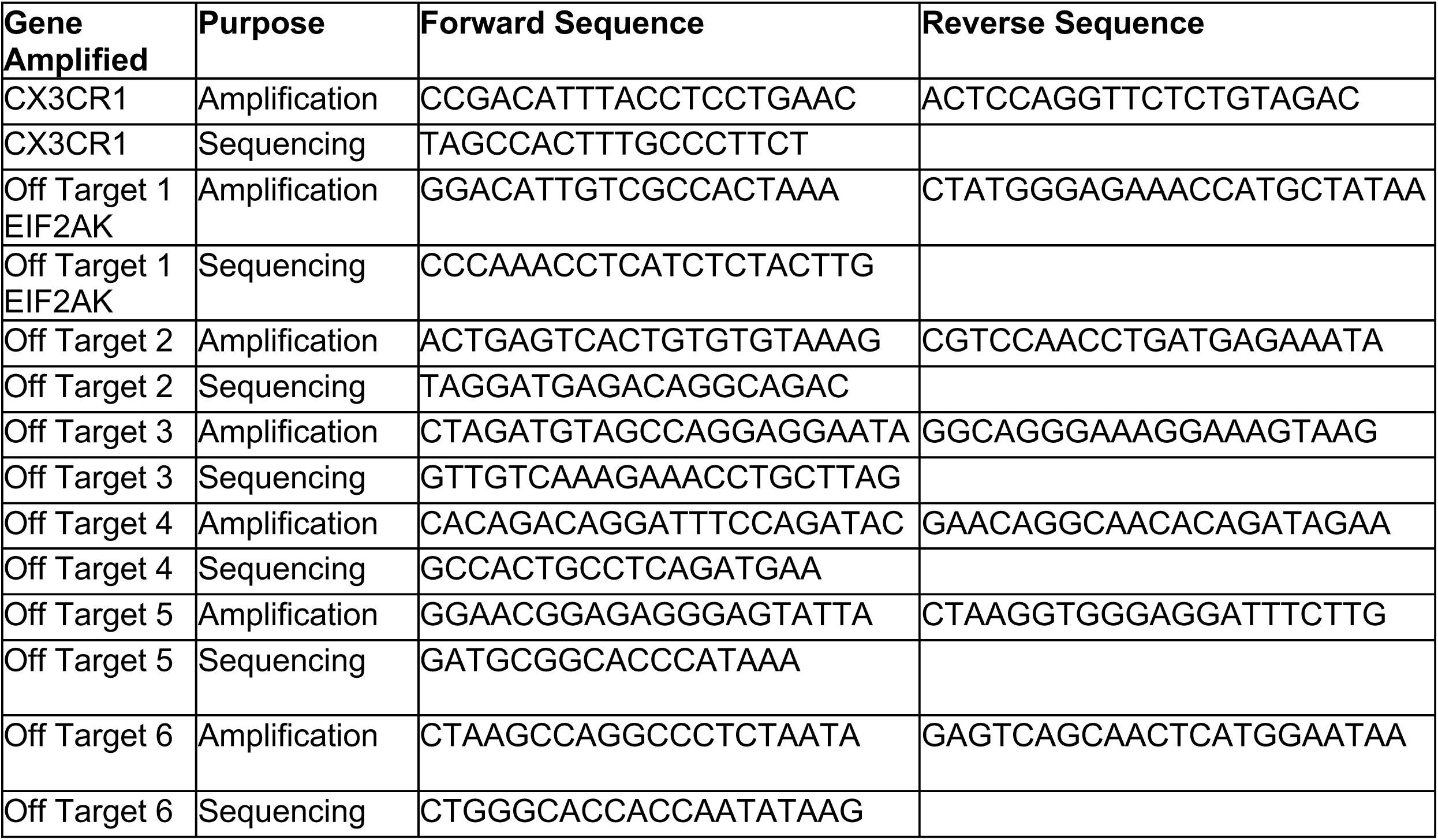
Primers used for PCR amplification and sequencing to confirm CRISPR/Cas9 editing and lack of off-target effects.

**Supplemental Figure 1:**
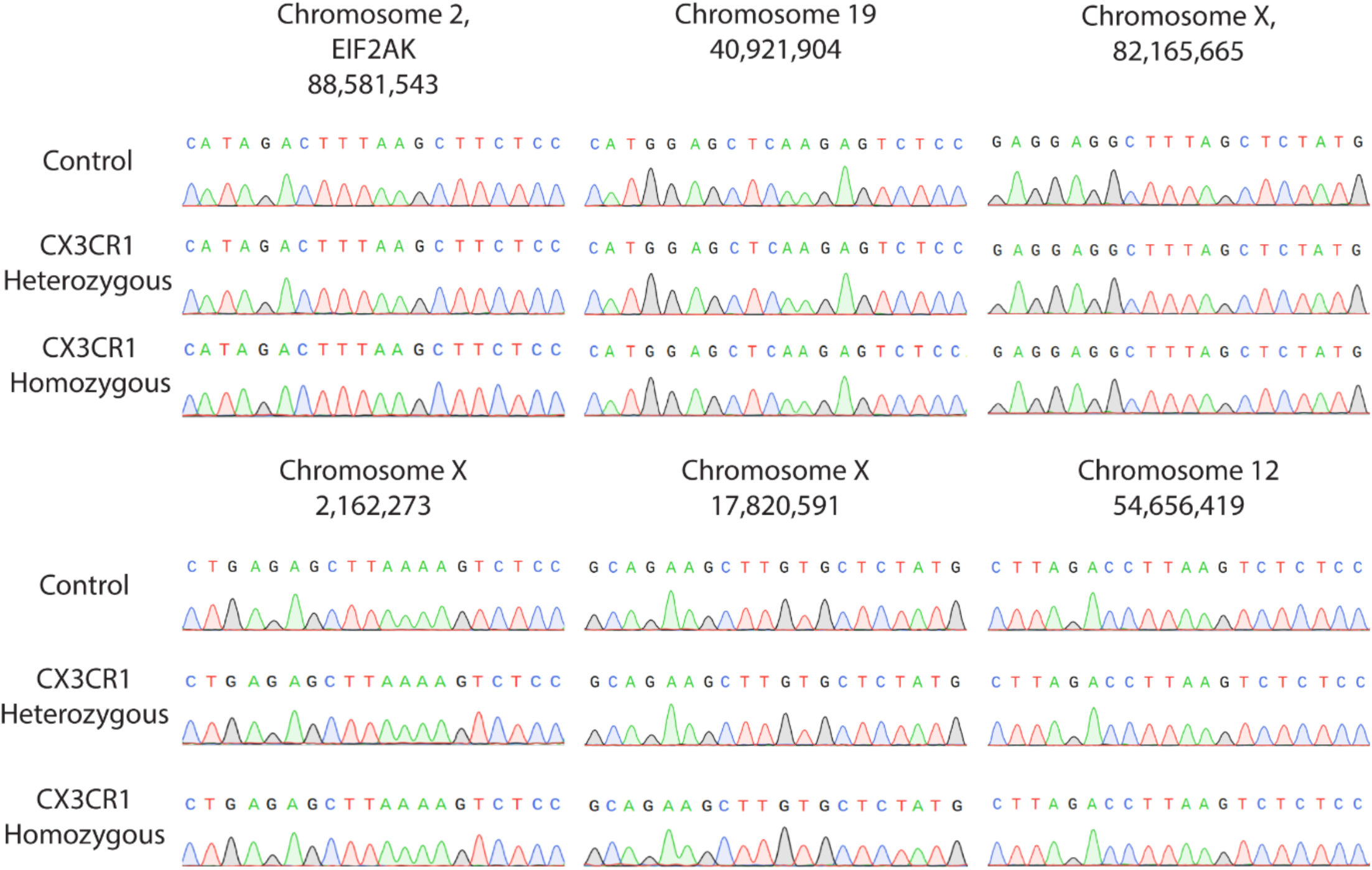
Sequencing of CX3CR1-V249I cell lines revealed no off-target effects. The top six most likely off target effects were sequenced in control, heterozygous V249I, and homozygous V249I cell lines and revealed no differences across genotypes due to off-target editing.

**Supplemental Figure 2:**
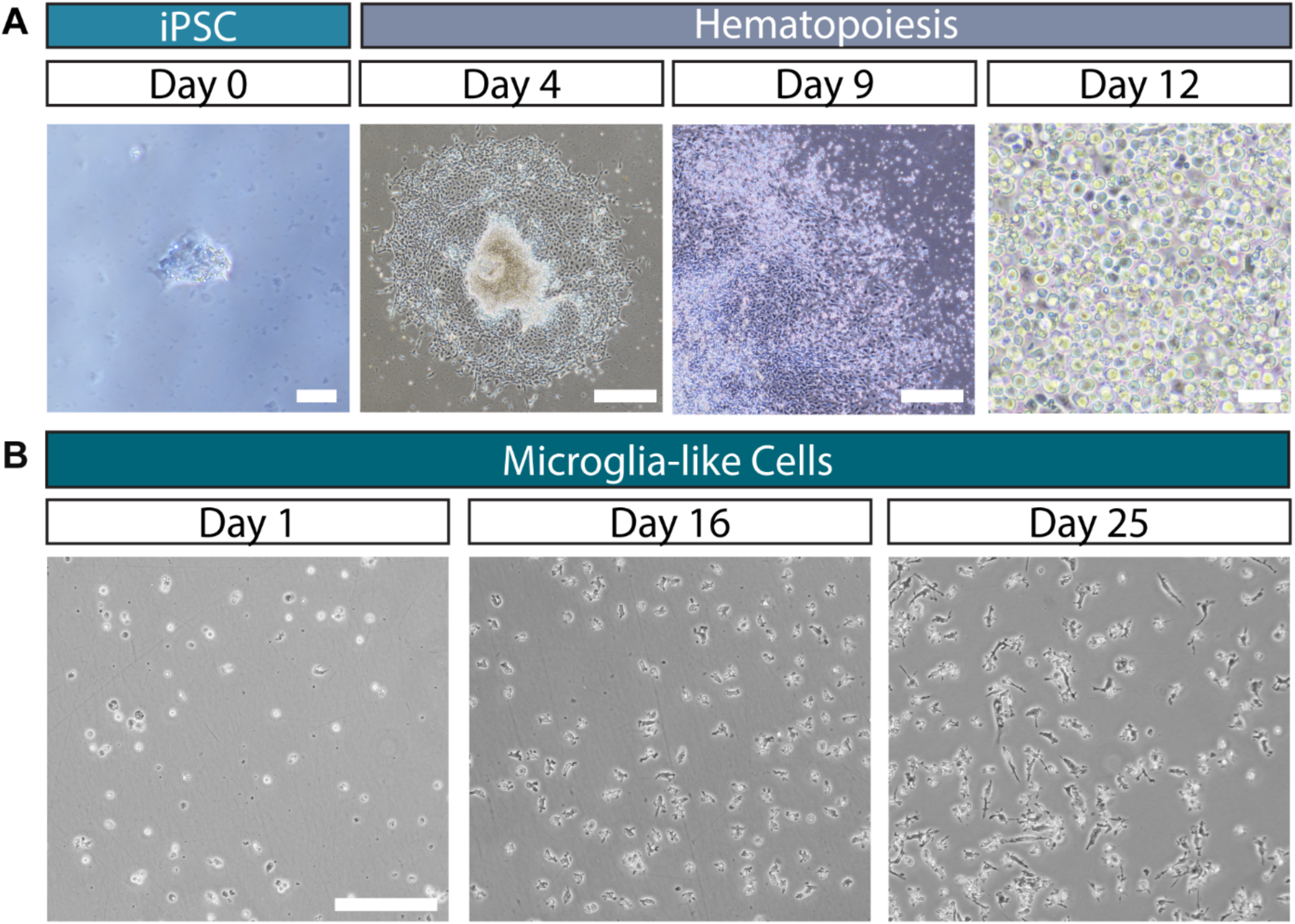
Differentiation of hMGLs. (A) Differentiation of iPSCs initially into hematopoietic progenitor cells. On day 0, iPSCs of approximately 150uM in size were attached. Scale: 100uM. On day 4, endothelial colonies began to form. Scale: 500uM. By day 9, endothelial colonies expanded in size and began producing HPCs. Scale: 500uM. On day 12, a large population of bright, round HPCs was visible and collected. Scale: 100uM. (B) Subsequent differentiation of HPCs into hMGLs using supplementation with mCSF, TGFb, and IL-34. Across 25 days of maturation, cells develop increasingly complex morphologies indicative of maturation. Scale: 500 µm.

**Supplemental Figure 3:**
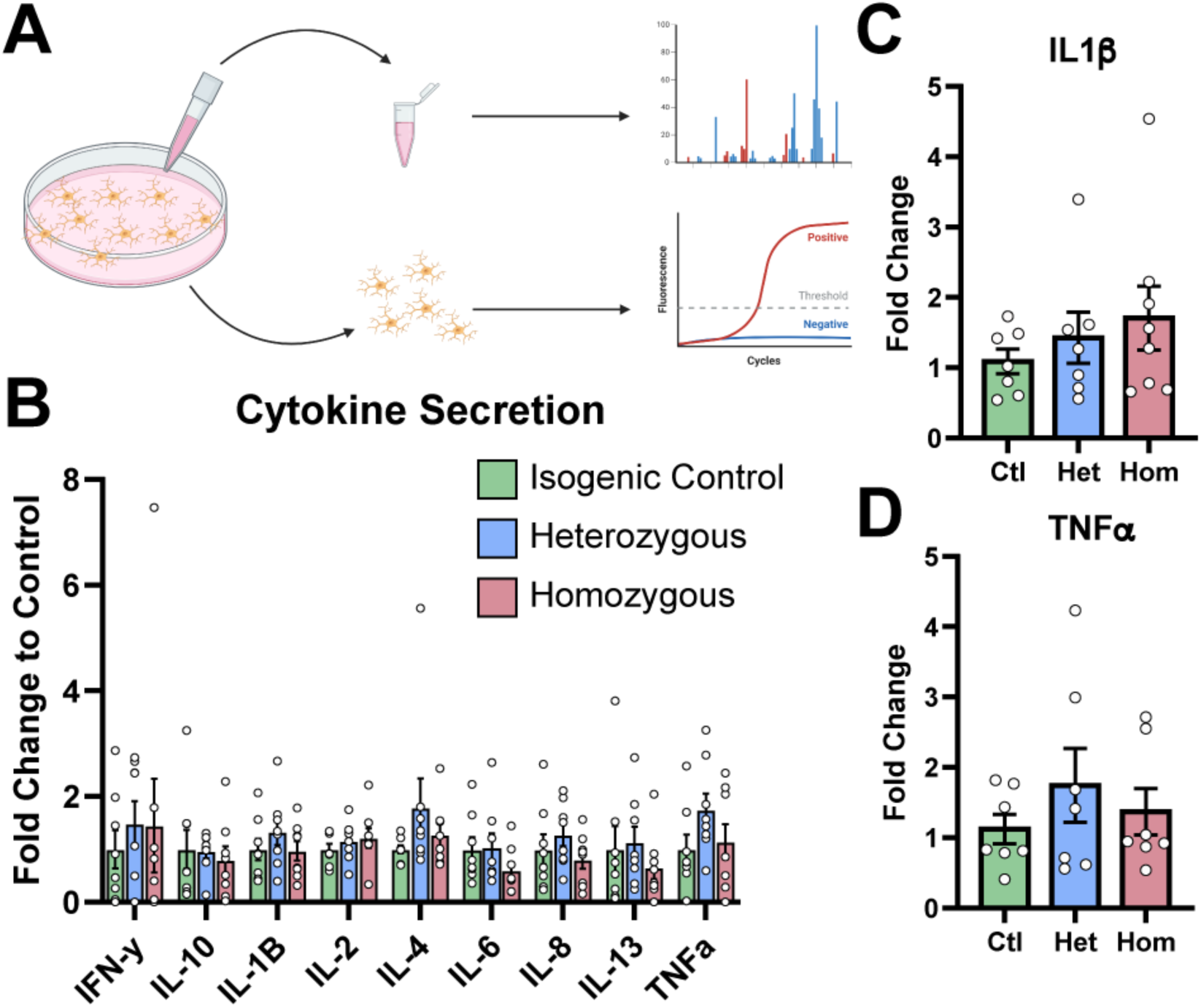
No differences in pro-inflammatory cytokine secretion were observed in V249I hMGLs. (A) Schematic representing collection of conditioned media from mature hMGLs for analysis and collection of RNA for qPCR. (B) The Mesoscale Discovery Kit V-PLEX Proinflammatory Panel 1 Human Kit was used to analyze conditioned media from hMGLs. No differences were observed in levels of any cytokine. (C-D) qPCR revealed no difference in IL1β or TNFα expression across hMGL genotype.

## Notes

### Competing Interest Statement

The authors have declared no competing interest.

